# The Zpr-3 antibody recognizes the 320-354 region of Rho and labels both rods and green cones in zebrafish

**DOI:** 10.1101/2022.02.21.481375

**Authors:** Pan Gao, Yayun Qin, Zhen Qu, Yuwen Huang, Xiliang Liu, Jingzhen Li, Fei Liu, Mugen Liu

## Abstract

Retinal markers with high quality and specificity are important for the observation of pathologic changes of retinal cells during retinal development, degeneration, and regeneration. Zpr-3 is a mouse monoclonal antibody with uncharacterized antigen usually used to label rod photoreceptors. In this study, we provided evidence to demonstrate that the antigen gene of zpr-3 is *rho*, which encodes the rod opsin, and the exact epitope of zpr-3 is the 320-354 region of Rho protein. More importantly, our immunofluorescence assays indicated that zpr-3 labels both the outer segments of rods and green-cones on zebrafish retinal sections, probably due to the cross-reaction with the green-cone opsin. Our work is valuable for the scientific community to interpret the experimental data involving the zpr-3 antibody.

## Introduction

Zebrafish has emerged as a useful animal model for research on retinal diseases in recent years ^1, 2^. The laminated structure of the retina is well-conserved between zebrafish and other vertebrates ^3^, and most pathogenic genes of inherited retinal degeneration have orthologues in zebrafish. Additionally, zebrafish retina shows strong regeneration ability ^4-6^. These make zebrafish a powerful model to study retinal development, degeneration, and regeneration. Antibodies labeling the five types of photoreceptors are important tools for investigating the changes of specific photoreceptors in Zebrafish ^7, 8^. The zpr-3 antibody is developed by ZIRC (http://zfin.org/ZDB-ATB-081002-45), and is widely used to label rods in zebrafish ^9-11^. However, the exact antigen of zpr-3 is still unknown.

In this study, we found that the band pattern of zpr-3 is quite similar to the zebrafish Rho antibody in western blot assays. Further experiments strongly indicated that the zpr-3 antigen is the rod opsin. Besides, immunofluorescence assays showed that in addition to rods’ outer segments, zpr-3 antibody also recognizes the outer segments of green-cones on zebrafish retinal sections.

## Methods

### Cell culture and zebrafish culture

ARPE-19 (ATCC Cat# CRL-2302) cells were authenticated by STR analysis and cultured in DMEM:F12 medium (Gibco) containing 10% FBS in a humid CO_2_ incubator described previously^12^. The full-length and truncated zebrafish *rho* were cloned into the p3xFlag CMV7.1 vector. ARPE-19 cells were transfected with the constructed plasmids respectively using the Lipofectamine 3000 reagent (Invitrogen) according to the manual. Wild-type and Tg(rho:EGFP) (a gift from Dr. Jian Zhou, Zhejiang University) zebrafish were cultured at 28.5 °C in a recirculating water system with a 14-h light/10-h dark daily cycle described previously^13^. All procedures were approved by the Ethics Committee of Huazhong University of Science and Technology.

### Western blot and immunofluorescence assays

For western blot assays, ARPE-19 cells were washed twice with PBS and lysed with the RIPA buffer containing protease inhibitor cocktail (Sigma, P8340) 24 hours after transfection. Zebrafish were sacrificed by immersing in 0.02% MS222 (Sigma, E10505) until no opercular movement. Eyes were enucleated and sonicated in RIPA lysis. The protein lysates were mixed with loading buffer and incubated at 95 °C for 5 minutes. Western blot was performed as previously described^14^. The following primary antibodies were used: zpr-3 (ZIRC, 1:250), anti-Rho antibody (customized, 1:500), anti-Flag antibody (MBL, M185, 1:5000), anti-GFP antibody (Abmart, M20004, 1:5000).

For immunofluorescence assay, ARPE-19 cells were fixed in 4% PFA in PBS for 15 minutes. Zebrafish eyes were fixed with 4% PFA overnight, cryo-protected in 30% sucrose, embedded in OCT, and sectioned along with the dorsal-ventral orientation through the optic nerve. The immunofluorescence was performed as previously described ^14^. The following primary antibodies were used: anti-Flag antibody (Abclonal, AE063, 1:1000), zpr-3 (ZIRC, 1:200), anti-Opn1lw antibody (customized, 1:100), anti-Opn1mw antibody (customized, 1:100).

## Results

### Zebrafish *rho* gene is the antigen gene of zpr-3

Firstly, we found that the band pattern of zpr-3 was exactly similar to the customized zebrafish Rho antibody in western blot assays of zebrafish retinal samples (Figure 1A). Next, Flag-tagged Rho protein was ectopically expressed in ARPE-19 cells and detected by western blot using anti-Flag, anti-Rho and zpr-3 antibodies, respectively. The three antibodies produced similar ladder-like bands (Figure 1B), demonstrating that Rho is likely the antigen of zpr-3. We also performed immunofluorescence assays in ARPE-19 cells. The overlap between red fluorescence signals of zpr-3 and green fluorescence signals of anti-Flag antibody again confirmed our western blot results (Figure 1C).

**Figure 1.**
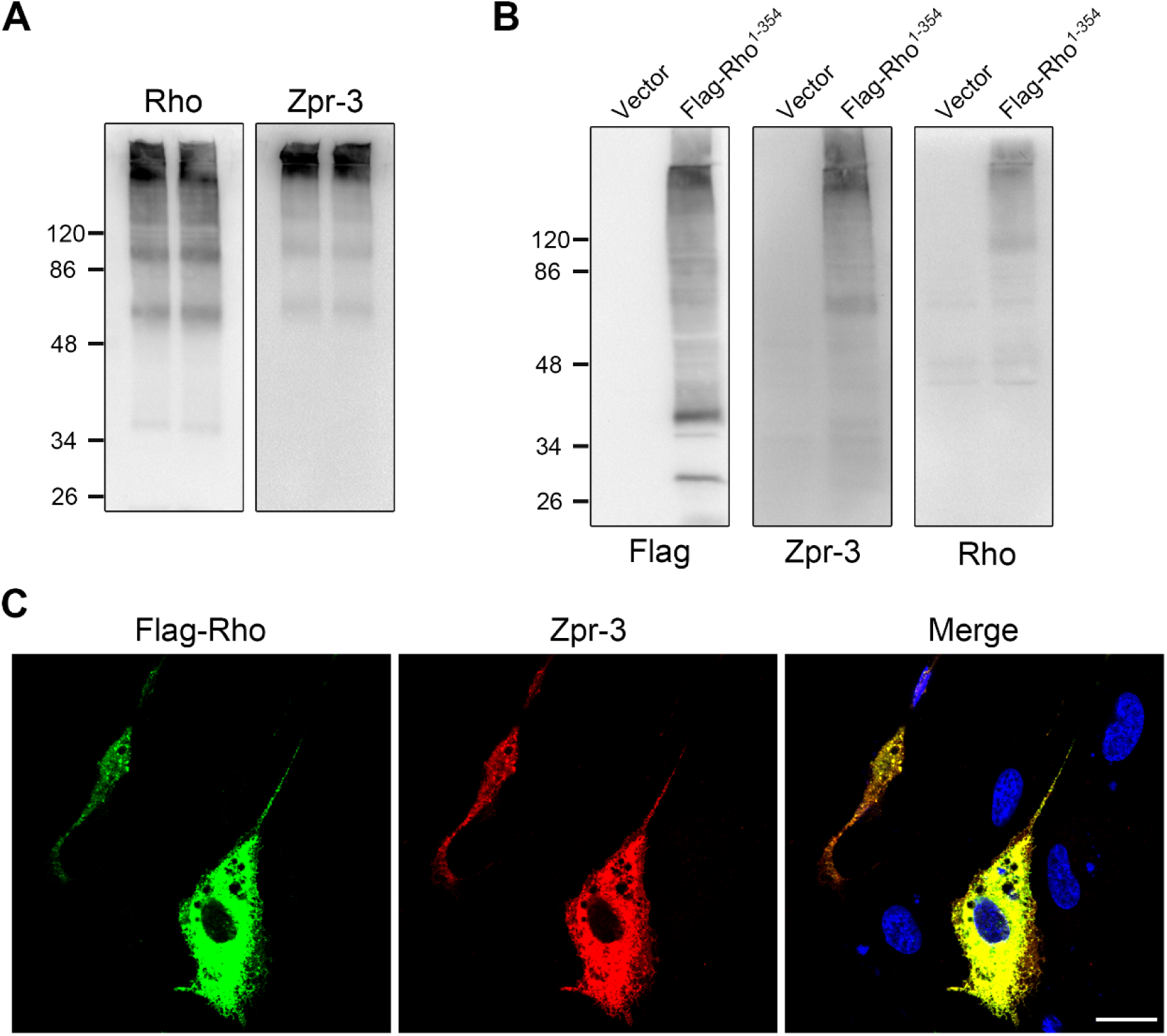
The antigen of zpr-3 is zebrafish Rho protein. (A) The ladder-like bands produced by zpr-3 and the anti-Rho antibody in the western blot assays of wild-type zebrafish protein lysates. (B) Full-length zebrafish Rho protein tagged with Flag was expressed in ARPE-19 cells and detected by western blot using the zpr-3, anti-Rho and anti-Flag antibodies, respectively. The images of western blot were shown. (C) Overlapping of the fluorescence signals of zpr-3 and anti-Flag antibody in ARPE-19 cells transfected with the zebrafish *rho* expression plasmid. Scale bar, 20 μm.

### The exact epitope of zpr-3 is the 320-354 region of Rho protein

To map the exact epitope of zpr-3, a series of truncated Rho mutants were constructed and expressed in ARPE-19 cells. Both zpr-3, and anti-Rho antibody reacted with the Rho fragments containing the C-terminal 320-354 region in western blot assays (Figure 2A-C). Further experiments showed the 338-354 region of Rho is sufficient for the recognition of anti-Rho antibody, while the 320-354 region is essential for zpr-3 recognition (Figure 2D). These results were summarized in a schematic diagram (Figure 2E). Our work indicated that the 320-354 region of Rho is the epitope of zpr-3.

**Figure 2.**
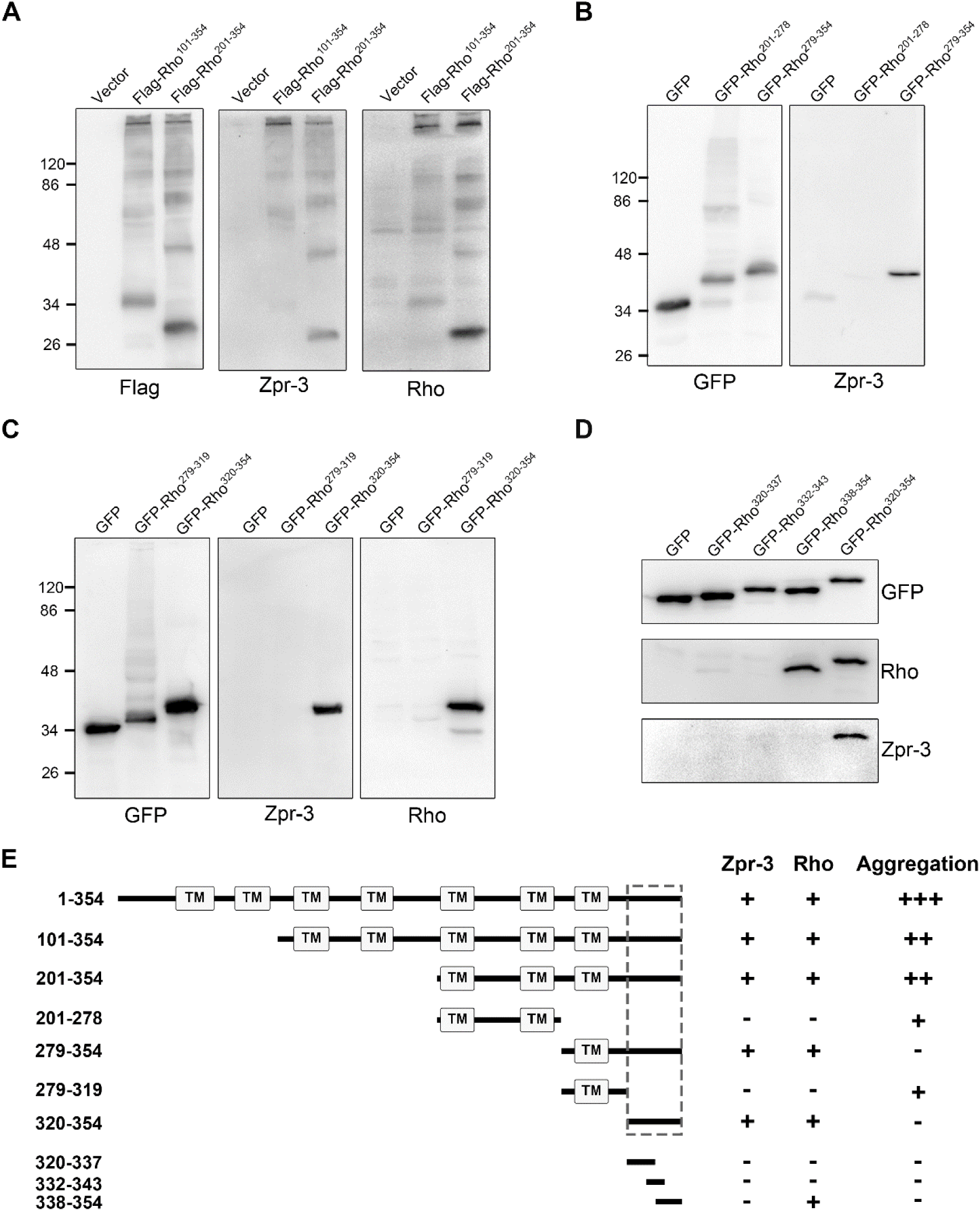
Identification of the exact epitope of zpr-3. Truncated zebrafish Rho proteins, 101-354 and 201-354 (A), 201-278 and 279-354 (B), 279-319 and 320-354 (C), 320-337, 332-343and 338-354 (D), were expressed in ARPE-19 cells and detected by western blot using the zpr-3 and anti-Rho antibodies, respectively. The images of western blot were shown. (E) The schematic summary of the above western blot results was shown. The multiple transmembrane domains of Rho were indicated, and the epitope of zpr-3 was highlighted by the dashed frame.

### Zpr-3 labels both rods and green-cones in zebrafish retinal sections

The zpr-3 antibody showed more fluorescence signals than the anti-Rho antibody in the immunofluorescence assays of zebrafish retinal sections at 14 dpf (Figure 3A). Using the Tg(rho:EGFP) transgene Zebrafish, we found that in addition to the EGFP labeled rods, zpr-3 also labeled the outer segments of other photoreceptors at 7dpf (Figure 3B).

**Figure 3.**
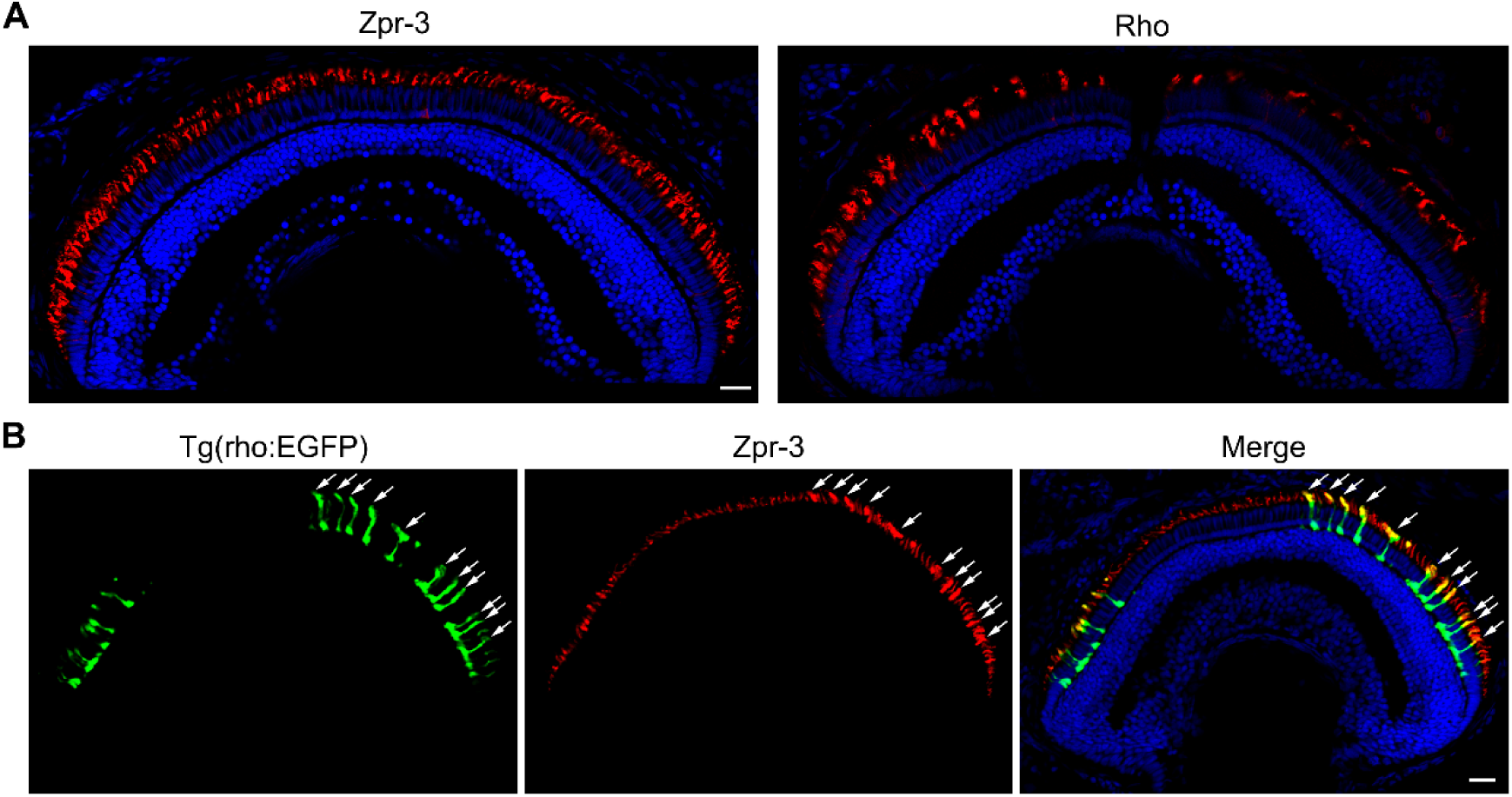
The fluorescence signals of zpr-3 are not solely located in rods. (A) Immunofluorescence results of zpr-3 and the anti-Rho antibody in 14 dpf zebrafish retinal sections. Scale bar, 20 μm. (B) Partial co-localization of the red fluorescence signals of zpr-3 with the EGFP-labeled rods (arrows) in 7 dpf zebrafish retinal sections. Scale bar, 20 μm.

To confirm which types of cones contribute to the additional fluorescence signals, we performed double labeling using zpr-3 and the anti-Opn1lw (red-cone opsin) or anti-Opn1mw (green-cone opsin) antibodies. Zpr-3 labeled the outer segments of green-cones, but not red-cones (Figure 4A, 4B). The sequence alignment of the zpr-3 epitope with Opn1mw and Opn1lw showed that the 320-354 region of Rho is moderately similar to Opn1mw, but very different from Opn1lw, which might explain why zpr-3 cross-reacts with Opn1mw but not Opn1lw.

**Figure 4.**
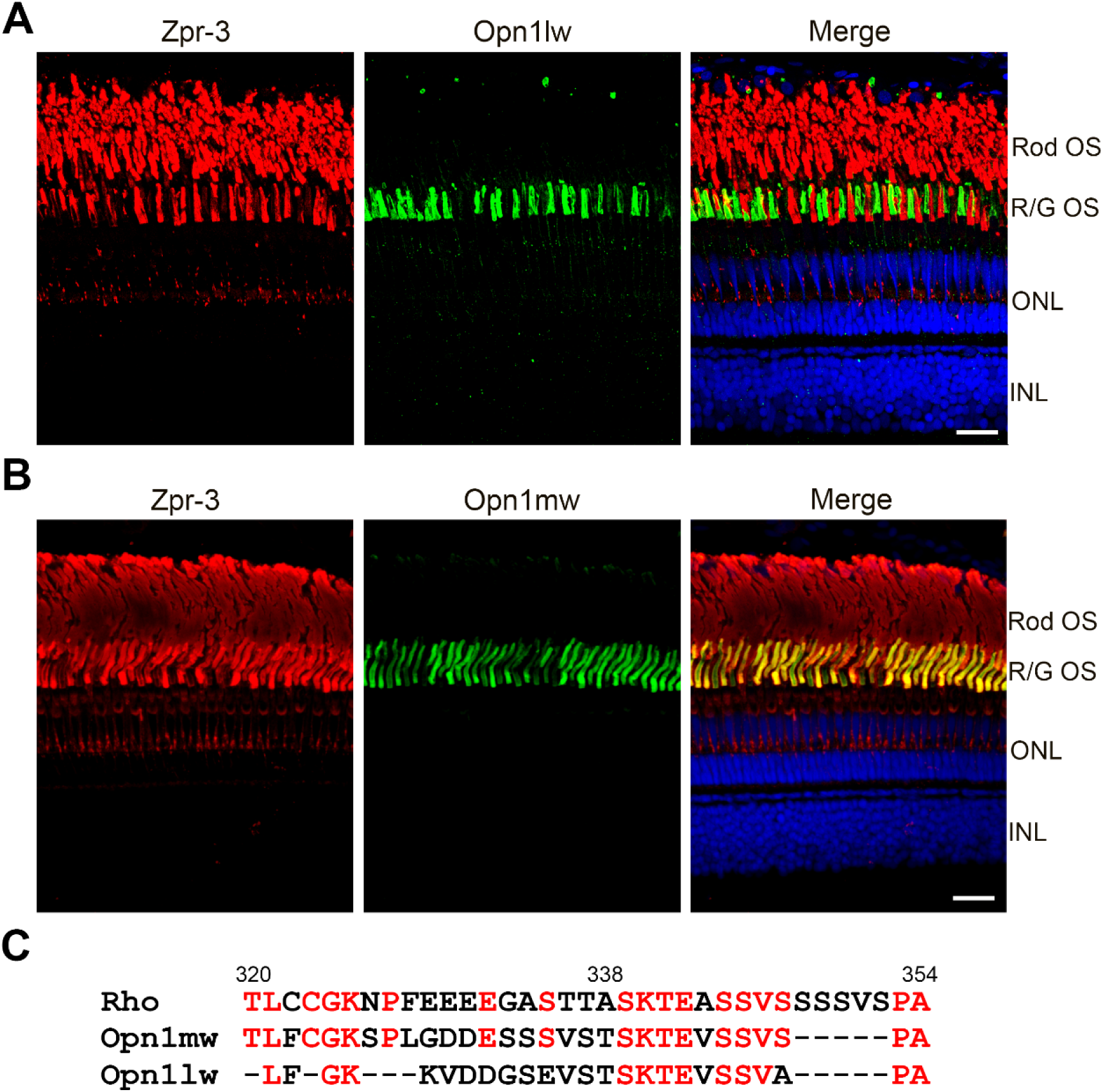
Zpr-3 labels green-cones as well as rods in immunofluorescence assays. (A) Double-label immunofluorescence images of zpr-3 and anti-Opn1lw antibody on zebrafish retinal sections. The fluorescence signals of zpr-3 were almost not co-localized with the outer segments of red-cones. Scale bar, 20 μm. (B) Double-label immunofluorescence images of zpr-3 and anti-Opn1mw antibody on zebrafish retinal sections. Obvious co-localization between the fluorescence signals of zpr-3 and the outer segments of green-cones was shown. Scale bar, 20 μm. (C) Sequence alignment of the epitope of zpr-3 (320-354 aa in Rho) and its corresponding regions in Opn1mw and Opn1lw. The identical amino acid residues were marked as red.

## Discussion

Zpr-3 is a widely used mouse monoclonal antibody. In this study, we provided evidence to demonstrate that the antigen gene of zpr-3 is *rho* and the exact epitope of zpr-3 is the 320-354 aa of Rho protein. More importantly, we demonstrated that zpr-3 labels the outer segments of both rods and green-cones in immunofluorescence assays, likely due to the cross-reaction with green-cone opsin. As an alternative marker for zebrafish rods, zpr-3 has been reported to stain double cone outer segments previously^8^. In this study, we further strengthened and extended the findings. Our results showed that the outer segments of rods and green-cones could be distinguished in adult zebrafish according to their locations. However, for zebrafish within 2 weeks of birth, it’s impossible to distinguish the two types of photoreceptors only based on the signals of zpr-3. In summary, our work will help the researchers to collect and interpret the data related to zpr-3 more accurately.

## Acknowledgments

We thank Dr. Jian Zhou (Zhejiang University) for providing the Tg(rho:EGFP) zebrafish. This work was supported by the National Natural Science Foundation of China [31601026, 31571303, 81670890, 31801041 and 81800870].

